# Zika virus oncolytic signaling reverts immunosuppressive and angiogenic glioblastoma microenvironment *in vitro* and *in vivo*

**DOI:** 10.64898/2026.09.20.752714

**Authors:** Gabriella P. A. de Freitas, Jefferson H. Quintanilla, José M. Janeiro, Oleksandra Brechko, Romy Weinstock, Gabriel C. Atella, Matheus V. Andrade, Jessica C.C.G. Ferreira, Janaina M. de Vasconcelos, João Vianez, Maria Bellio, Amilcar Tanuri, Luiz G. Dubois, Flávia R.S Lima, Vivaldo Moura-Neto, Luiza M. Higa, Patricia P. Garcez

**Author notes:** Correspondence to L.M.H and P.P.G. These authors contributed equally to this work (Luiza M. Higa, Patricia P. Garcez).

## Abstract

Glioblastoma (GBM), the most aggressive primary brain tumor, remains refractory to conventional therapies, characterized by high relapse rates and an extremely poor clinical prognosis. Oncolytic viruses represent a promising therapeutic frontier; specifically, Zika virus (ZIKV) has shown potent oncolytic activity against GBM in vitro and in vivo. However, its impact on the tumor microenvironment (TME) remains poorly understood. Here, we demonstrate that ZIKV modulates pivotal TME components, exerting a paracrine anti-angiogenic effect that impairs endothelial stabilization and reprogramming microglia toward an anti-tumoral phenotype. These findings were corroborated in vivo in GBM-bearing mice. Using unbiased RNA-Seq, we identified type III interferon (IFN) signaling as the primary mediator of this ZIKV-induced TME remodeling. Crucially, analysis of human glioma datasets reveals that the type III IFN receptor complex is highly expressed in patients. This identifies a high-risk population that possesses the molecular machinery to respond to this signaling axis. Exogenous treatment with type III IFN *in vitro* and *in vivo* successfully mimicked ZIKV’s anti-tumoral effects, reducing angiogenesis and promoting immune activation. Our results demonstrate that ZIKV acts as a powerful immunotherapeutic agent capable of shifting the TME toward an anti-tumoral state. By targeting a signaling axis characterized by high receptor density in treatment-refractory cases, this study identifies the type III IFN pathway as a therapeutic vulnerability and a promising strategy for patients who have failed conventional therapies.

## INTRODUCTION

Glioblastoma (GBM) is the most aggressive and common primary tumor of the central nervous system (CNS) (*1*). Despite standard treatment, which includes surgical removal of the tumor mass, radiotherapy, and chemotherapy, the average survival of diagnosed patients is approximately 15 months, with only 10% of patients reaching the 5-year mark post-diagnosis (*2–4*). GBM exhibits a highly heterogeneous tumor microenvironment (TME), which is considered one of the factors responsible for treatment resistance (*5, 6*). The TME consists of cancer cells, neurons, endothelial cells, astrocytes, microglia, infiltrating immune system cells, and extracellular matrix components (*7–9*).

As GBM is a tumor that shows limited responsiveness to standard treatment, new therapeutic approaches are being studied and developed targeting tumor cells and its TME. In this context, one alternative is the use of oncolytic viruses (OVs), a therapy that employs unmodified or modified/attenuated replication-competent viruses that selectively infect and kill tumor cells without causing harm to the healthy cells surrounding the tumor (*10*). The lysis of tumor cells results in the release of tumor antigens, which stimulates the immune response against the tumor (*10*).

A new potential OV that has been studied in the context of GBM is the Zika virus (ZIKV). ZIKV is a flavivirus that was first isolated in 1947 in Uganda (*11*). In 2015, ZIKV reemerged and was associated with microcephaly and other CNS abnormalities in fetuses whose mothers were infected during pregnancy (*12–14*). ZIKV infection in adults is predominantly mild or asymptomatic and cases affecting the central nervous system are rare (*15, 16*). Interestingly, multiple studies have described the ZIKV oncolytic ability. In vitro studies using 3D models and GBM cell lines have demonstrated the strong oncolytic capacity of ZIKV, which is effective not only against GBM but also other CNS tumors, such as medulloblastomas and atypical teratoid rhabdoid tumors (*17, 18*).

In vivo, ZIKV administration in GBM-bearing mice increases survival, reduces tumor mass, and modulates immune cells positively (*17, 19*). Dogs with spontaneous CNS tumors showed no adverse effects from ZIKV treatment and exhibited reduced tumor size, extended survival, and improved neurological symptoms (*20*). In humans, a case report described a 43-year-old woman diagnosed with GBM and ZIKV infection shortly after surgery. Notably, she remained relapse-free for at least six years (*21*).

The tropism of ZIKV for GBM cells and GBM stem cells has been well demonstrated (*17, 22*). However, the effects of ZIKV infection on the GBM microenvironment are unknown. In the context of the GBM TME, endothelial cells promote significant tumor angiogenesis, while tumor-associated microglial cells secrete factors that stimulate tumor mass growth. Therefore, ZIKV’s tropism for GBM cells and the potential consequences of this infection in the TME offer the prospect of identifying new cellular and molecular targets for GBM treatment through a therapeutic strategy based on viral infection. Here we demonstrate the effect of ZIKV infection on the microglia and endothelial cells in the context of GBM in vitro and in vivo.

## RESULTS

### ZIKV infection selectively reduces glioblastoma cell viability via apoptosis

To evaluate the impact of ZIKV on glioblastoma (GBM) survival, we challenged four cell lines with varying MOIs (0.2 to 5) over a 7-day period. Based on their sensitivity to viral infection, cell lines were classified into two distinct groups: responsive and non-responsive, using a 50% viability reduction threshold. Under these criteria, GBM02 and T98G were characterized as non-responsive, as they maintained stable viability levels (Figure. 1, A and B). In contrast, sensitive lines exhibited robust cell death; TG1 cells reached the 50% threshold by 7 dpi (Figure. 1C), while U-87 MG showed a more rapid response, achieving a 70% reduction by 3 dpi (Figure. 1D).

**Figure 1.**
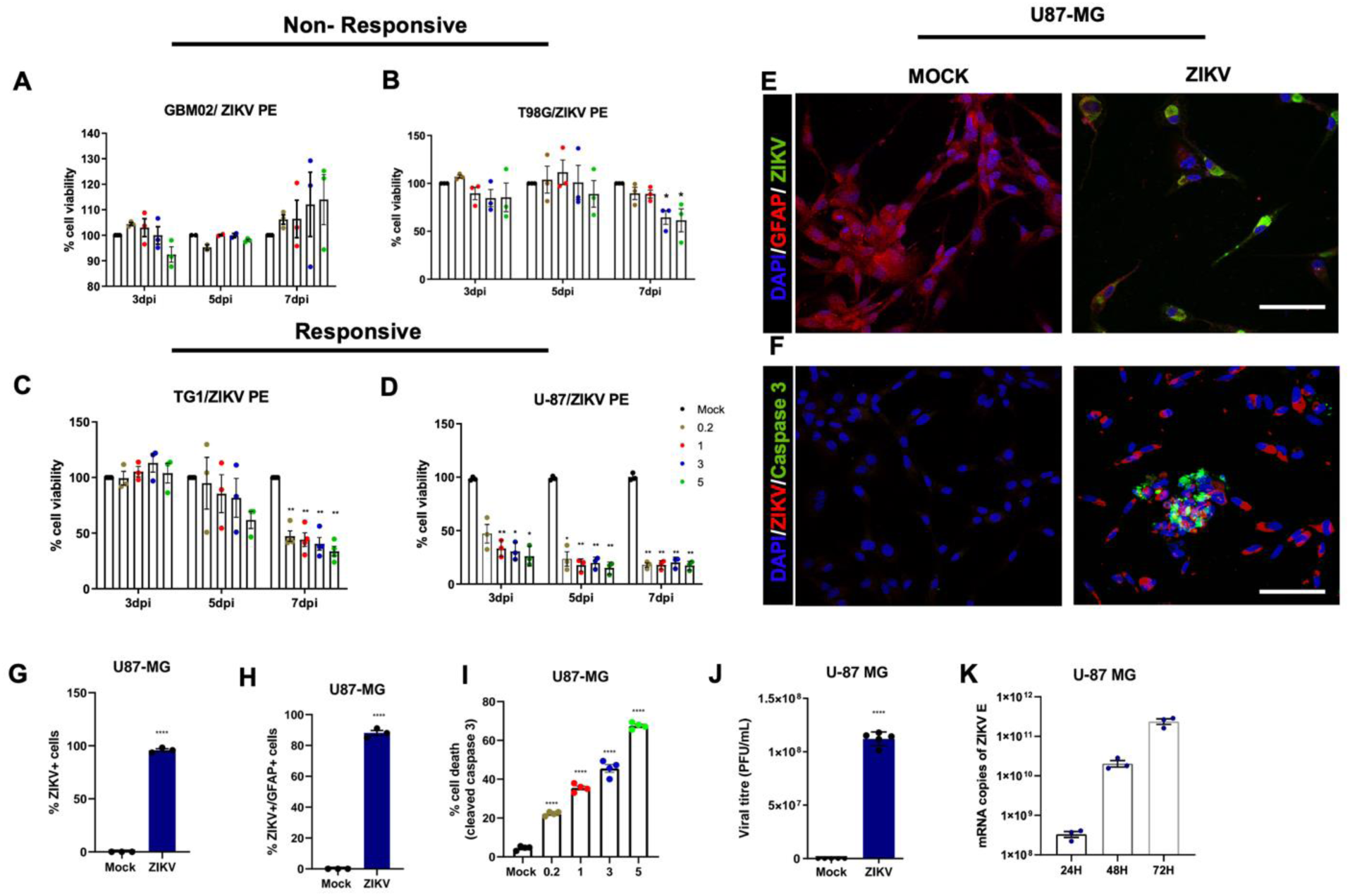
ZIKV infection selectively reduces glioblastoma cell viability via apoptosis. (**A** and **B**) Cell viability of non-responsive GBM02 and T98G cells upon ZIKV infection (MOIs 0.2, 1, 3, and 5) or mock treatment at 3, 5, and 7 days post-infection (dpi) (n=4). (**C** and **D**) Cell viability of responsive TG1 and U-87 MG cells under the same conditions (n=4). Responsiveness was defined as a viability reduction of at least 50%. (**E**) Representative immunofluorescence (IF) of U-87 MG cells at 3 dpi (MOI 3) showing ZIKV 4G2 (green) and GFAP (red). (**F**) Representative IF of ZIKV-infected U-87 MG cells at 3 dpi showing cleaved caspase-3 (green) and ZIKV (red). (**G**) Quantification of ZIKV-positive U-87 MG cells at 3 dpi (n=3). (**H**) Quantification of ZIKV-infected cells expressing the glial marker GFAP (n=3). (**I**) Quantitative analysis of cell death by counting cleaved caspase-3 positive cells at 3 dpi (n=3). (**J**) Quantification of infectious particles produced by ZIKV-infected U-87 MG cells at 3 dpi (n=3). (**K**) ZIKV mRNA viral load in U-87 MG cells at 3 dpi (n=3). Data are presented as mean ± SEM. *p<0.05, **p<0.01, ***p<0.001. Scale bar, 30 μm.

To characterize the infection, we performed immunofluorescence on U-87 MG cells, which revealed widespread expression of the viral marker 4G2 (green) alongside the astrocytic marker GFAP (red) (Figure. 1E). This reduction in viability was driven by programmed cell death, as evidenced by the expression of cleaved caspase-3 (Figure. 1F). Quantitative analysis confirmed high infection rates, with over 95% of cells being 4G2-positive in responsive and non-responsive cell line (Figure. 1G, Figure S1 A). Phenotypic characterization further showed that ZIKV infected cells expressing the glial marker GFAP (Figure. 1H). Consistently, we observed a significant increase in the percentage of apoptotic cleaved caspase-3 positive cells (Figure. 1I). Finally, we confirmed that both responsive and non-responsive lines supported productive viral replication, as demonstrated by the release of infectious particles (Figure. 1J, Figure S1 B) and a high viral mRNA load (Figure. 1K).

### ZIKV-infected glioblastoma cells exert paracrine anti-angiogenic effects

GBM cells secrete factors that modulate the tumor microenvironment (TME) and tumor growth. To investigate how ZIKV-infected GBM cells impact the TME, we utilized a paracrine model using conditioned medium (CM) from uninfected (CM-MOCK) or ZIKV-infected (CM-ZIKV) U-87 MG cells to treat non-tumoral components, specifically endothelial and microglial cells.

Endothelial cells are vital for maintaining vascular integrity and supporting tumor-driven angiogenesis. Cells treated with CM-ZIKV exhibited a significant reduction in migration at 48 hours, covering approximately 60% less area compared to CM-MOCK controls (Figure. 2, A and B). Furthermore, CM-ZIKV treatment led to a decrease in endothelial cell proliferation by approximately 20% (Figure. 2C). To assess endothelial barrier function, we analyzed the expression of the tight junction protein ZO-1. Our results revealed a significant reduction in ZO-1 expression accompanied by a disorganized distribution pattern in cells treated with CM-ZIKV, indicating a compromised capacity to form an effective endothelial barrier (Figure. 2, D and E).

**Figure 2:**
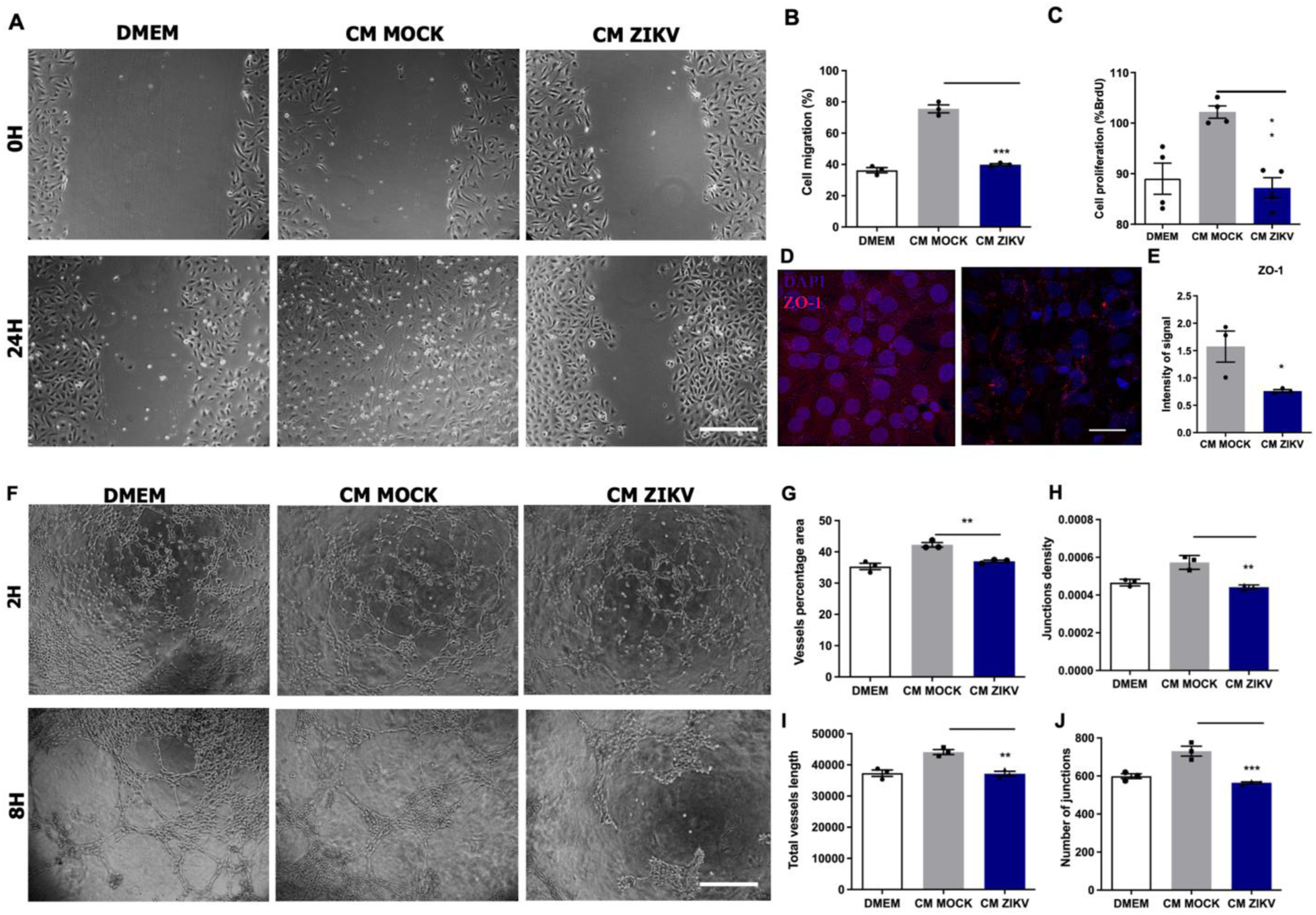
ZIKV-infected GBM secretome impairs endothelial function and angiogenesis *in vitro*. **(A and B)** Representative wound healing assay images of HBMECs treated with CM-MOCK or CM-ZIKV at 0 and 24 hours post-treatment, with corresponding quantification of cell migration (n=3). **(C)** Assessment of endothelial cell proliferation via BrdU incorporation after 24 hours of treatment with CM-MOCK or CM-ZIKV (n=4). **(D and E)** Representative immunofluorescence (IF) and quantitative analysis of the tight junction marker ZO-1 (red) in HBMECs after 24 hours of treatment, showing reduced intensity and pattern disorganization in the CM-ZIKV group (n=3). **(F)** Representative images of Matrigel tube formation assay after 8 hours of treatment. **(G-J)** Quantitative assessment of angiogenic parameters, including (**G**) percentage of vessel area, (**H**) junction density, (**I**) average vessel length, and (**J**) total number of junctions (n=3). Data are presented as mean ± SEM. *p<0.05, **p<0.01, ***p<0.001. Scale bar, 30 μm.

To further explore these anti-angiogenic effects *in vitro,* we conducted a Matrigel tube formation assay. CM-ZIKV markedly disrupted the formation of capillary-like structures at 8 hours compared to the CM-MOCK condition (Figure. 2F). This disruption was characterized by a significant decrease in vessel area (Figure. 2G), lower junction density (Figure. 2H), and shorter average tube length (Figure. 2I) and fewer overall junctions (Figure 2J).

To validate these findings in vivo, we performed the Matrigel plug assay in adult mice. Matrigel plugs implanted with CM-ZIKV appeared less dense and more translucent compared to those with CM-MOCK, with markedly fewer visible blood foci (Figure 3A). Histological analysis revealed that CM-MOCK plugs exhibited robust neovascularization, characterized by the presence of vessel-like structures containing red blood cells (black arrows Figure 3B, F), consistent with effective angiogenesis. In contrast, plugs from the CM-ZIKV group showed a pronounced reduction in neovascular structures formation and contained significantly fewer red blood cell-filled structures (Figure 3C, F). Furthermore, we found a substantial decrease in the number of endothelial cells within the CM-ZIKV plugs (Figure 3D, G) compared to CM-MOCK (Figure 3E, G), suggesting impaired endothelial cell recruitment and/or proliferation in response to factors secreted by ZIKV-infected GBM cells. Collectively, these findings indicate that ZIKV infection not only hampers endothelial cell proliferation and migration but also severely compromises their ability to participate effectively in the complex process of angiogenesis.

**Figure 3:**
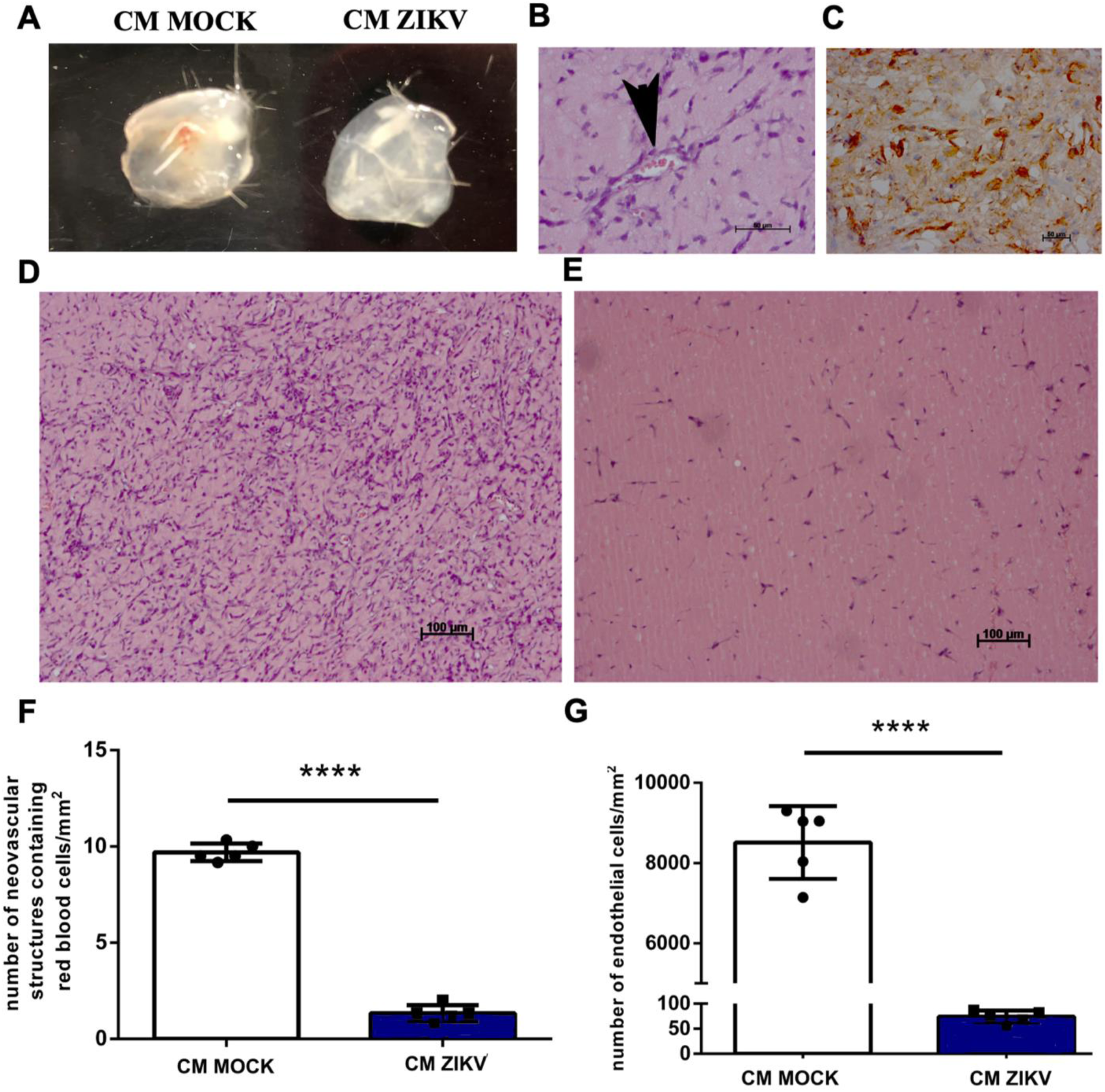
ZIKV-infected GBM secretome impairs angiogenesis *in vivo*. **(A)** Representative macroscopic images of Matrigel plugs harvested from adult mice. CM-MOCK plugs (left) exhibit a dense, vascularized appearance with visible blood foci, whereas CM-ZIKV plugs (right) are translucent and less vascularized. **(B and C)** Representative histological sections showing neovascular structures containing red blood cells (black arrowheads) in CM-MOCK (B) versus the pronounced reduction of these structures in CM-ZIKV (C) plugs. **(D and E)** Hematoxylin and eosin (H&E) staining showing a marked reduction in the total number of endothelial cells in CM-ZIKV (E) compared to CM-MOCK (D) plugs. **(F)** Quantitative analysis of neovascular structures per section (n=6 plugs per group). **(G)** Quantification of the total number of endothelial cells per field (n=6 plugs per group). Data are presented as mean ± SEM. ****p<0.0001. Scale bars, 50 and 100 μm.

### ZIKV infection reprograms microglia toward an anti-tumoral phenotype

Microglial cells, as resident immune cells of the central nervous system, play a crucial role in shaping the GBM TME. To investigate how ZIKV infection in GBM affects the functional state of microglia, primary microglia cultures were treated with CM ZIKV. Our findings revealed that microglia treated with CM-ZIKV exhibited significantly higher levels of inducible nitric oxide synthase (iNOS) compared to CM-MOCK controls (Figure. 4, A to G). This upregulation of iNOS is a hallmark of microglial activation toward a pro-inflammatory, anti-tumoral state. In addition to increased iNOS expression, we observed a significant rise in microglial proliferation following exposure to CM-ZIKV (Figure. 4H).

**Figure 4:**
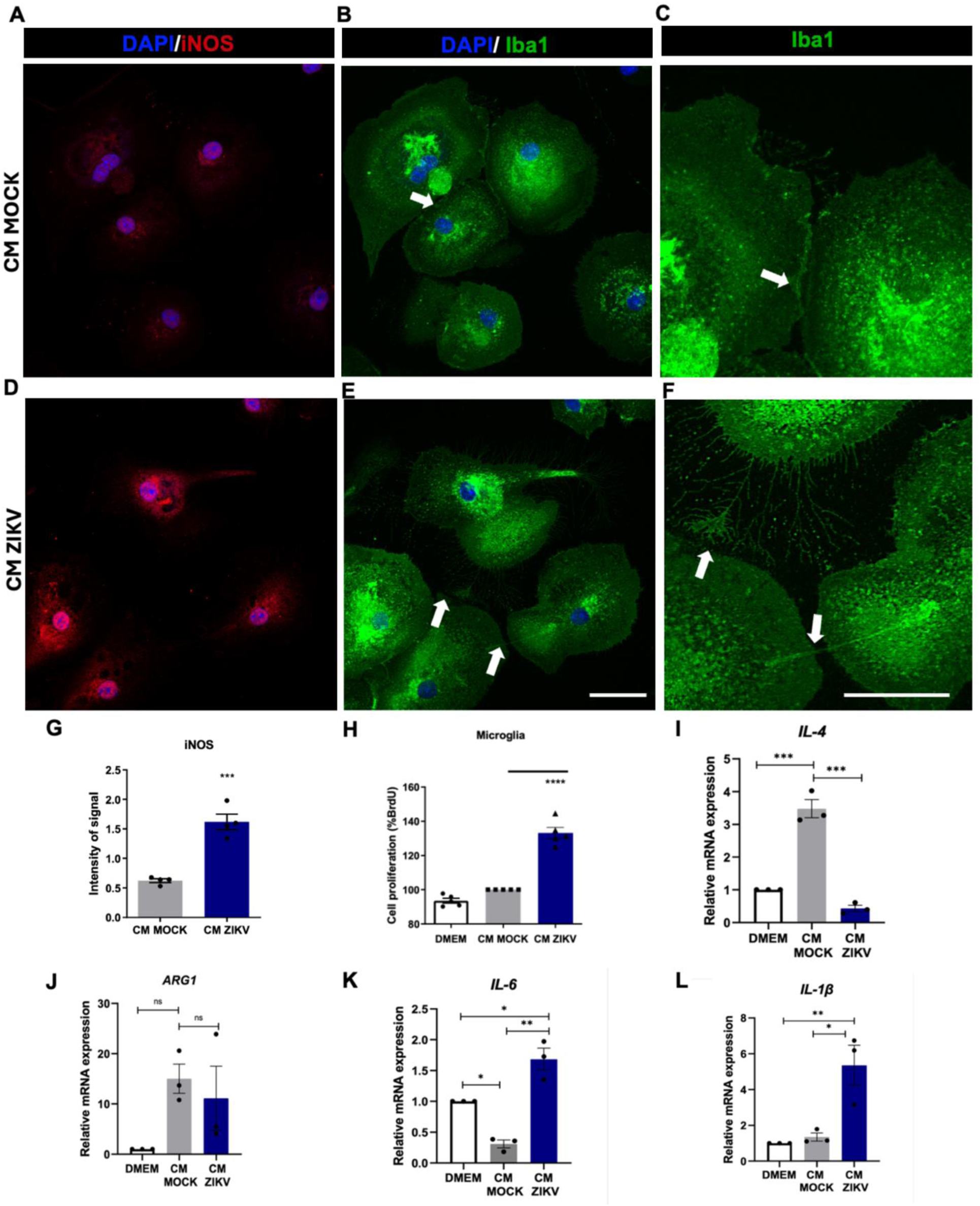
Reprogramming of microglia following exposure to conditioned media (CM) of Zika virus (ZIKV)-infected GBM cells. **(A-C)** Representative immunofluorescence (IF) of microglia treated with CM-MOCK showing (**A**) iNOS (red), (**B**) Iba1 (green), and (**C**) a magnified view highlighting the lack of cellular ramifications (white arrow). **(D-F)** Representative IF of microglia treated with CM-ZIKV showing (**D**) increased iNOS expression (red), (**E**) Iba1 expression (green), and (**F**) a magnified view highlighting abundant cellular ramifications (white arrows), indicative of an activated state. **(G)** Quantitative analysis of iNOS expression levels (n=3). **(H)** Assessment of microglial proliferation 48 hours post-treatment (n=3). **(I-L)** Cytokine expression profile of microglia treated with CM-MOCK or CM-ZIKV, including **(I**) IL-4, (**J**) ARG1, (**K**) IL-6, and (**L**) IL-1β (n=3). Data demonstrate a shift from a pro-tumoral (ARG1/IL-4) to an anti-tumoral (IL-6/ IL-1β) profile. Data are presented as mean ± SEM. *p<0.05, **p<0.01, ***p<0.001. Scale bar, 10 μm.

Typically, the TME in GBM is characterized by pro-tumoral microglia and macrophages that contribute to local immunosuppression. These immune cells facilitate immune evasion by tumor cells and promote tumor progression enhancing tumor survival and invasion (*28, 29, 30*).

Furthermore, ZIKV infection induced a distinct shift in the cytokine expression profile of microglia. Specifically, we observed a marked reduction in the expression of IL-4 (Figure. 4I) and arginase 1 (Fig. 4J), both of which are classical markers associated with pro-tumoral, immunosuppressive microglial polarization. Conversely, exposure to the ZIKV-infected secretome triggered an increase in the expression of the pro-inflammatory cytokines IL-6 (Figure. 4K) and IL-1β (Figure 4L). Collectively, these results suggest that ZIKV infection reprograms the TME by diminishing pro-tumoral characteristics and enhancing the capacity of microglia to mount an effective anti-tumor immune response.

### GBM Tumor is reduced in ZIKV-infected mice

To investigate the impact of ZIKV infection on the GBM microenvironment, we employed an established *in vivo* mouse glioma model. This model involved the intracranial injection of GL261 mouse glioma cells into wild type Swiss mice. Seven days post-inoculation, mice received a single intratumoral injection of ZIKV (Figure 5A). While tumors were visible from the cortical surface in both groups after 7 days of infection, MOCK-treated tumors (Figure 5B) were markedly larger than ZIKV-infected tumors (Figure. 5C).

**Figure 5.**
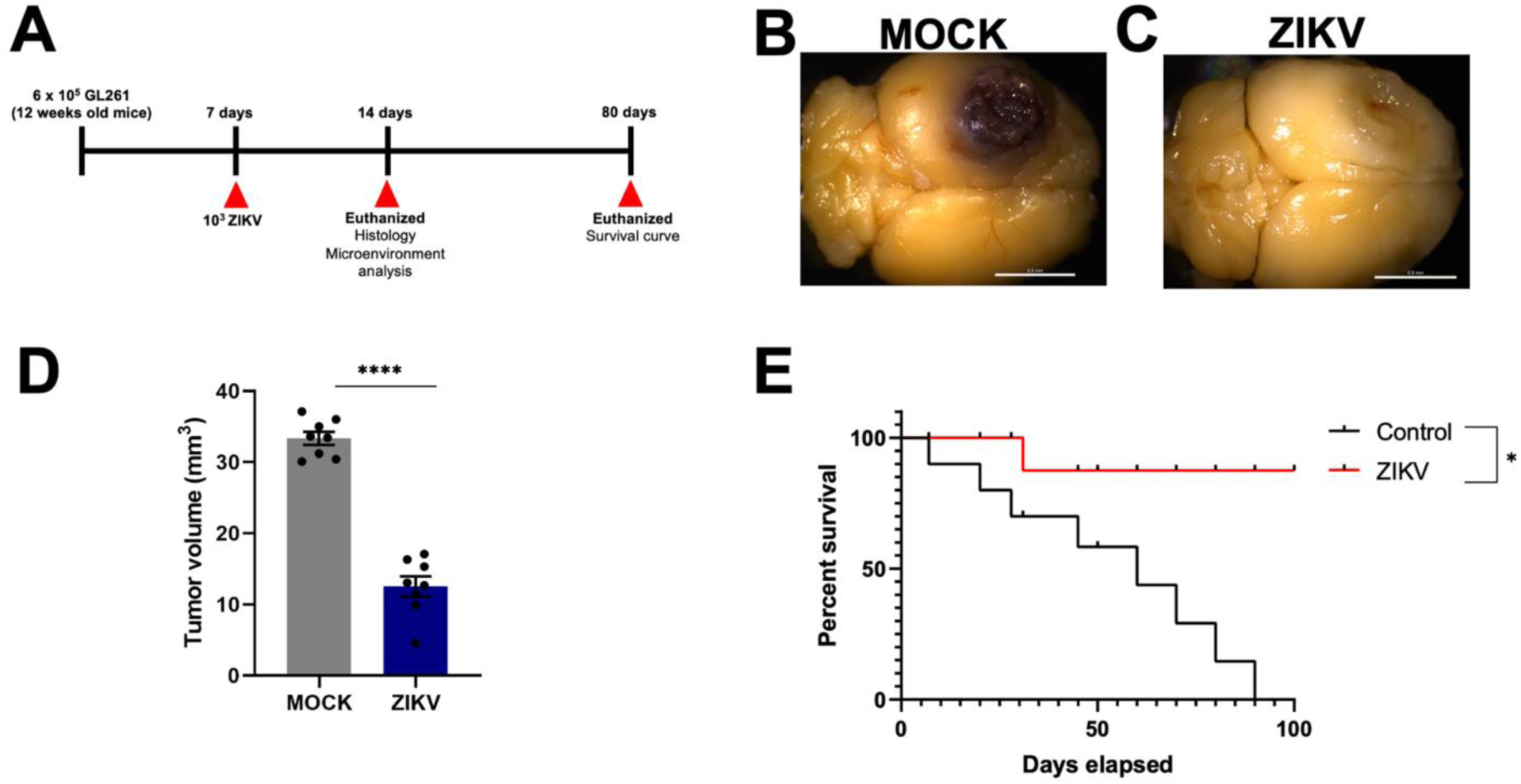
Intratumoral ZIKV infection restricts glioblastoma progression and extends survival *in vivo*. **(A)** Schematic representation of the experimental timeline, illustrating GL261 cell inoculation, intratumoral ZIKV or MOCK treatment, and subsequent analysis. **(B and C)** Representative whole-brain images from mice at 7 days post-infection showing visibly larger tumors in the MOCK group (B) compared to ZIKV-infected mice (C). **(D)** Stereological quantification of tumor volume based on histological slice analysis (n=8 per group). **(E)** Kaplan-Meier survival curves showing significantly extended lifespan in ZIKV-treated mice compared to the MOCK-treated group (n=8 per group). Statistical significance was determined using the Log-rank test. Data are presented as mean ± SEM. *p<0.05, ****p<0.0001$. Scale bar, 0.5 mm.

This suggests that ZIKV infection may restrict tumor expansion, a finding that contrasts with the typical rapid growth observed in untreated GBM. Quantitative analysis of tumor size across multiple histological slices corroborated our initial observations, with ZIKV-infected tumors displaying a significantly smaller volume compared to their MOCK counterparts (Figure 5F). This reduction in tumor volume is indicative of ZIKV’s capacity to impair GBM growth, an effect that was further validated by survival analysis. Mice bearing ZIKV-infected tumors demonstrated an extended lifespan relative to those in the MOCK-treated group (Figure 5G). Collectively, these findings demonstrate that ZIKV infection exerts a potent anti-tumoral effect in vivo, likely driven by a combination of direct viral cytotoxicity and therapeutic modulation of the tumor microenvironment.

### ZIKV suppresses tumor angiogenesis and enhances microglial infiltration *in vivo*

Given that angiogenesis is a critical process in the development and progression of malignant gliomas, driving tumor growth and expansion, we sought to investigate the effects of ZIKV infection on tumor vascularization. To this end, we quantified the density of IB4+ cells in GBM tissue slices. Notably, ZIKV infection and viral persistence were confirmed in the tumor tissue from day 7 (Supplementary figure 2A) up to 30 days post-infection (Supplementary figure 2B). Our analysis revealed a significant reduction in IB4 expression in ZIKV-infected GBM samples compared to the MOCK-treated controls (Figure 6A, B), indicating a notable decrease in the overall vasculature within the tumor microenvironment. Further analysis of angiogenic parameters provided additional insights into the impact of ZIKV on tumor angiogenesis. Specifically, we observed a marked decrease in the density of vessel junctions (Figure 6C) and a corresponding reduction in the total number of these junctions (Figure 6D), which are critical for the formation of complex vascular networks. Additionally, vessels in the GBM MOCK condition were found to be significantly larger and more extensive than those in the ZIKV-infected tumors (Figure 6E), leading to a significant reduction in the area occupied by vessels in the ZIKV-infected group (Figure 6F). These findings strongly indicate that ZIKV infection in GBM not only reduces the density and complexity of tumor vasculature but also impairs crucial angiogenic processes essential for tumor growth and progression.

**Figure 6.**
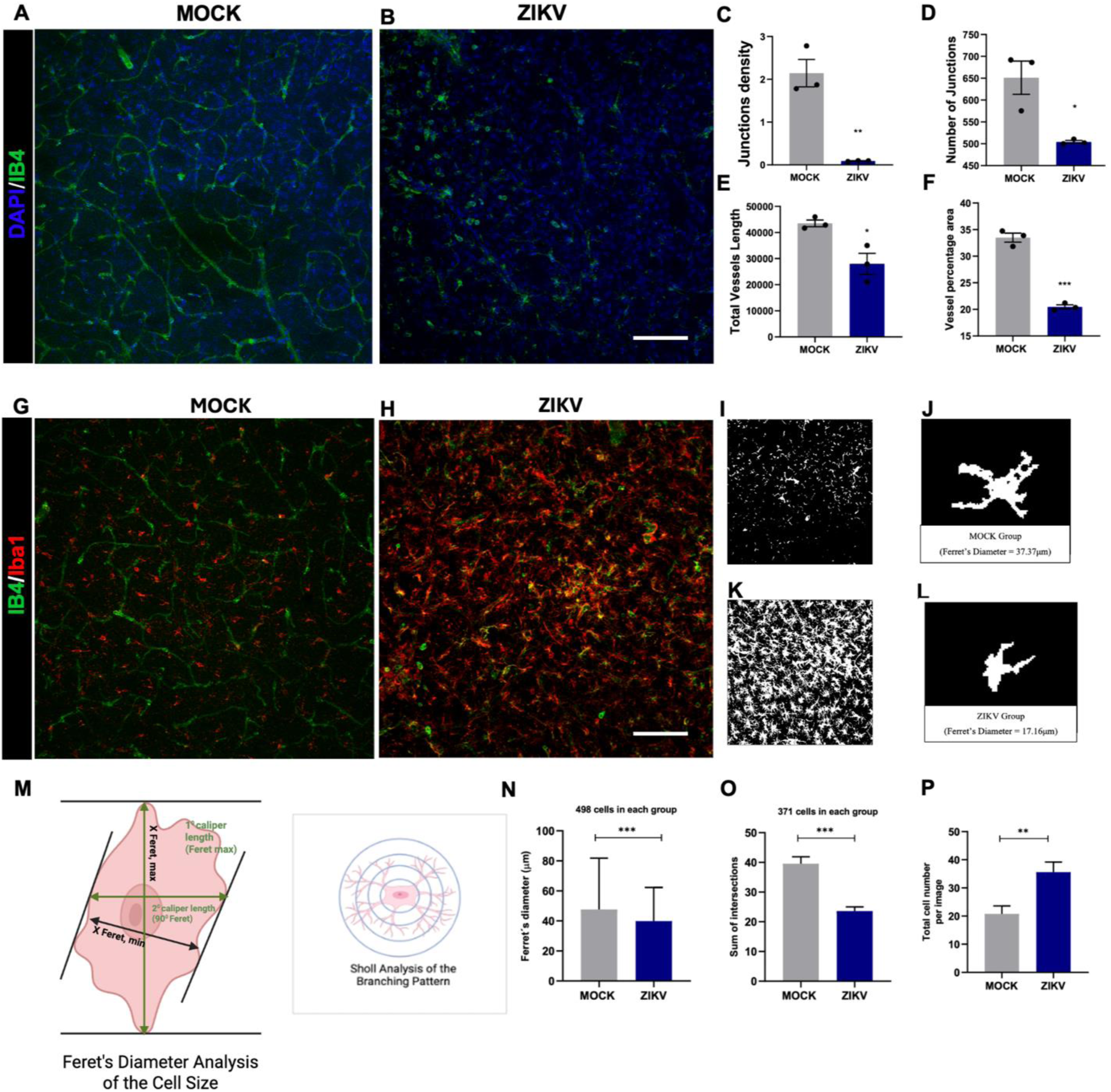
ZIKV infection impairs tumor vascularization and promotes microglial activation *in vivo*. **(A and B)** Representative immunofluorescence (IF) images of tumor blood vessels (IB4, red) in MOCK-treated (A) and ZIKV-infected (B) mice. **(C to F)** Quantitative assessment of vascular parameters, including (C) vessel junction density, (D) total number of junctions, (E) average vessel length, and (F) total area occupied by vessels (n=3 per group). **(G and H)** Representative IF images and quantitative analysis of microglial density (Iba1, red) in MOCK (G) versus ZIKV-infected (H) tumors. **(I to L)** Representative high-magnification images illustrating microglial morphology in MOCK (I, J) and ZIKV-infected (K, L) groups. **(M)** Schematic representation of Ferret’s diameter and Sholl analysis used for morphological characterization. **(N to P)** Quantitative morphological analysis showing (N) Ferret’s diameter (cell size, n=498 cells), (O) sum of intersections from Sholl analysis (ramification, n=371 cells), and (P) total microglial cell number. Data demonstrate a shift toward an activated phenotype with reduced size and ramification. Data are presented as mean ± SEM. *p<0.05, **p<0.01, \*\*\**p*<0.001. Scale bar, 30 *μ*m.

We observed that ZIKV infection in GBM-bearing mice leads to a significant increase in the density of Iba1+ cells, suggesting a robust enhancement in microglial recruitment when compared to MOCK-treated animals (Figure 6G, H). Our *in vitro* findings provide further insight into this phenomenon, showing that, indirectly, ZIKV promotes a shift in microglial activation, potentially priming microglia towards a pro-inflammatory state. The direct influence of ZIKV on microglia results in increased recruitment and notable morphological changes, when we compare MOCK-treated (Figure 6I, J) with ZIKV-treated (Figure 6K, L) mice. In addition to increased infiltration, ZIKV-exposed microglia exhibited distinct morphological changes characteristic of an activated, pro-inflammatory state (Figure 6M). This combined approach allowed us to evaluate changes in the ramification, shape and size of microglial cells across both groups. Our analysis revealed that microglial cells in the ZIKV-treated group were 26% smaller than those in the MOCK group, with average cell diameters of 28.39μm for MOCK and 20.91μm for ZIKV-infected GBM mice (Figure 6N). Sholl analysis of microglia from ZIKV-infected GBM animals demonstrated that ZIKV exposure induces microglia to adopt a more amoeboid morphology, which sharply contrasts with the extensively branched morphology typical of the MOCK condition (Figure 6 J, L). Moreover, there was a significant reduction in microglial ramification in the ZIKV group (Figure 6O). This morphological transformation is characteristic of microglial activation, indicating a shift from an anti-inflammatory, pro-tumor state to a pro-inflammatory, anti-tumor state. The alteration in microglial morphology from a resting, branched form to an activated, amoeboid form suggests a fundamental change in microglia function.

Furthermore, the significant increase in the total number of microglial cells observed in the parenchyma of ZIKV-infected GBM animals (Figure 6P) further supports the concept of a ZIKV-induced activation leading to a pro-inflammatory and anti-tumor phenotype. This influx of activated microglia into the tumor microenvironment likely contributes to the overall reduction in tumor burden seen in ZIKV-infected GBM mice, as well as the associated improvements in survival outcomes.

### ZIKV infection activates a type III interferon signaling axis

Given that ZIKV infection *in vivo* modulates the tumor-associated endothelium and microglial landscape, we sought to elucidate the molecular underpinnings of these effects. Previous studies have demonstrated that ZIKV challenges in neuro-oncological models trigger a robust activation of the innate immune response, characterized by the upregulation of type I, II, and type III IFN pathways (17, 31, 32, 33). Consistent with these reports, our transcriptomic analysis of ZIKV-infected GBM cells revealed a profound reprogramming of the transcriptional landscape (Figure. **7**A). Specifically, we identified a signature heavily enriched for antiviral and inflammatory mediators, with a prominent induction of the type III IFN signaling pathway.

**Figure 7.**
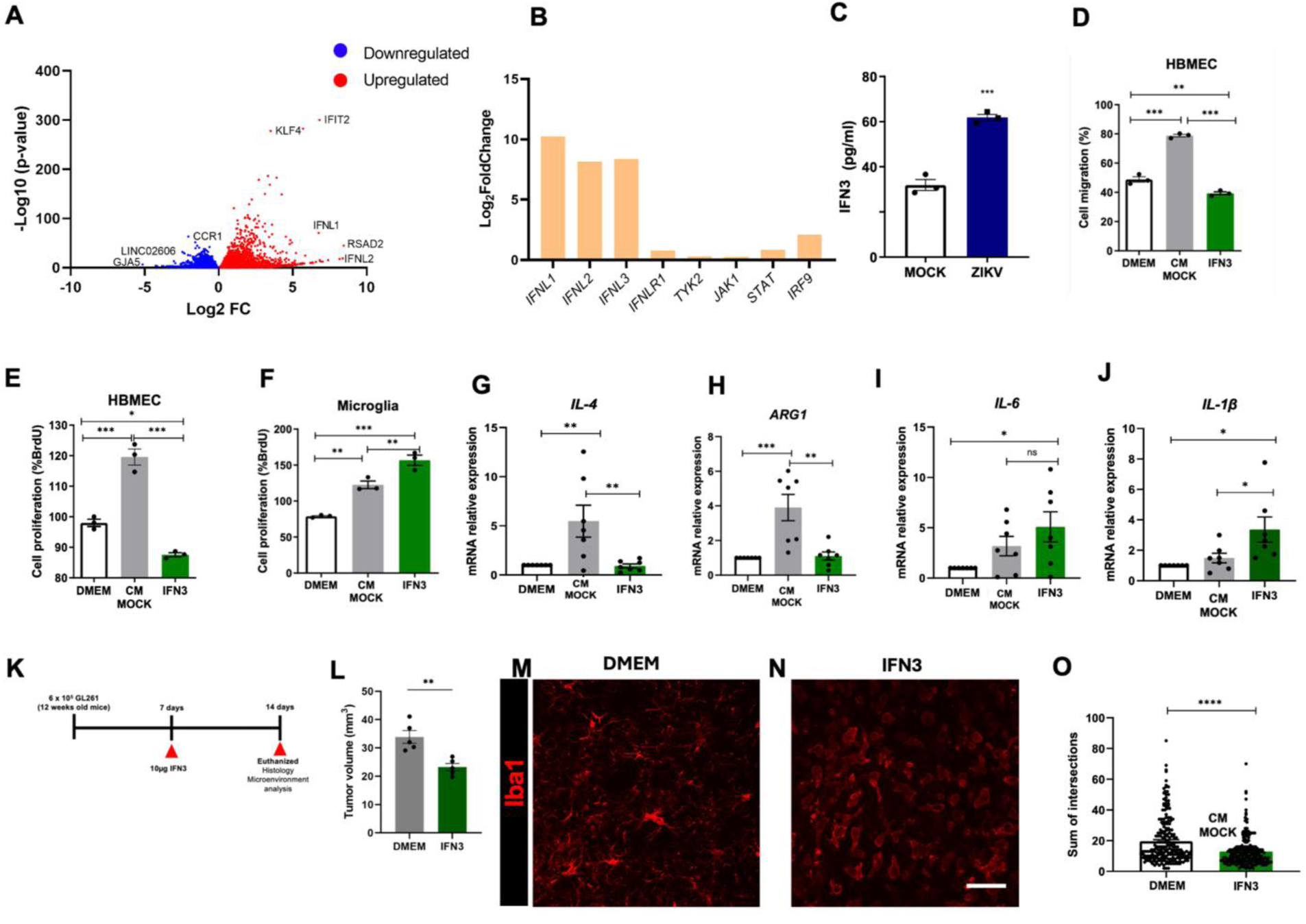
ZIKV triggers a type III interferon axis that modulates GBM microenvironment. (**A**) Volcano Plot showing the magnitude and statistical significance of gene expression changes following ZIKV infection, Genes significantly downregulated (blue) and upregulated (red). (**B**) Differential gene expression (DGE) in ZIKV infected U-87 MG cells (**C**) Quantification of IFN3 in ZIKV CM by ELISA assay. (**D**) Quantification of endothelial cell migration (n=3). (**E**) Quantification of endothelial proliferation (n=3). (**F**) Microglial proliferation (n=3). (**G, H**) Relative expression levels of pro-tumoral cytokine Interleukin-4 (IL-4) (**G**) and Arginase-1 (ARG1) (H) (n=4). (**I**, **J**) Relative expression levels of anti-tumoral cytokines Interleukin-6 (IL-6) (I) and Interleukin-1β (IL-1β) (J) (n = 4) (**K**) Schematic experimental timeline and design, illustrating the treatment and analysis schedule. (**L**) Tumor volume following type III IFN treatment. (**M, N**) Representative immunofluorescence image of tumor-associated microglia treated with control media (DMEM) (**M**) or IFN3 (**N**). (**O**) Quantification of the sum of intersections from Sholl analysis indicating a decrease in microglial ramification in the type III-treated group. *p<0.05, **p<0.01, ****p<0.001

Among the differentially expressed genes (DEGs), we observed a marked upregulation of key effectors and transducers of this cascade, including *IFNL1, IFNL2,* and *IFNL3*, as well as their cognate receptor *IFNLR1* and downstream signaling components such as *TYK2, JAK1, STAT2,* and *IRF9* (Figure. 7B). Quantitatively, *IFNL1* exhibited a 10.2-fold increase in transcript levels in ZIKV-infected GBM cells (p < 0.001), while *IFNL2* and *IFNL3* showed approximately 8-fold higher expression compared to MOCK-treated controls (p < 0.001). These transcriptomic findings were further validated by RT-qPCR, which confirmed a significant increase in the mRNA levels of type III IFN in ZIKV-infected GBM cells (Supplementary Figure 3 A-D). This robust transcriptomic profile supports the hypothesis that ZIKV-infected GBM cells undergo a secretome reprogramming characterized by a potent type III IFN output, which may drive the anti-angiogenic and pro-inflammatory remodeling observed in the tumor microenvironment (TME).

To establish the clinical relevance of these observations, we investigated the expression of the type III IFN receptor subunits, *IFNLR1* and *IL10RB*, in human patients using the CGGA database. We observed that both receptor subunits are expressed across various glioma subtypes, with levels reaching their peak in GBM (Supplementary Figure 3E, F). Notably, in the recurrent glioma setting, high expression of these receptors correlated with shorter overall survival (Supplementary Figure 3G, H). This indicates that the type III IFN receptor complex is a hallmark of the most aggressive and treatment-resistant phenotypes. Crucially, the high availability of these molecular targets in recurrent cases reinforces the potential of type III IFN-based therapies as a viable strategy for patients who have exhausted standard treatment options.

The activation of this pathway suggests that type III IFN is a key mediator of the immunomodulatory and antitumor effects observed in the ZIKV-conditioned medium. To validate this at the protein level, we performed ELISA analysis of the CM from ZIKV-infected GBM cells, detecting a significant increase in the concentration of secreted type III IFN compared to MOCK controls (Figure 7C). To evaluate whether type III IFN orchestrates these paracrine alterations, we challenged distinct cellular components of the TME with exogenous type III IFN.

In endothelial cells, type III IFN treatment significantly impaired vascular dynamics, resulting in a 15% reduction in migratory capacity and a 20% decrease in cellular proliferation (Figure 7D, E). Parallel experiments with microglial cells recapitulated the phenotype induced by ZIKV-CM, characterized by increased proliferation rates (Figure 7F) and profound transcriptional remodeling. Specifically, type III IFN exposure led to the downregulation of the protumoral markers *IL-4* and *Arg1* (Figure 7G, H), while simultaneously upregulating the proinflammatory cytokines *IL-6* and *IL-1β* (Figure 7I, J).

To assess the therapeutic relevance of these findings *in vivo,* GBM-bearing wild-type mice received intratumoral administration of type III IFN one-week post-implantation (Figure 7K). This treatment led to a significant reduction in tumor volume compared to DMEM-treated controls (Figure 7L). Consistent with our *in vitro* data, the 7-day treatment regimen induced marked morphological shifts in the microglial population (Figure 7M, N). Microglia in type III IFN-treated animals transitioned from a ramified, homeostatic state to an amoeboid morphology with significantly reduced ramifications (Figure 7O), a hallmark of proinflammatory activation. Collectively, these findings identify type III IFN as a central mediator of ZIKV-induced TME remodeling and highlight its potential as a therapeutic strategy to overcome the immunosuppressive landscape of GBM.

## DISCUSSION

Here, we provide a comprehensive characterization of how Zika virus (ZIKV) infection orchestrates a fundamental reprogramming of the glioblastoma (GBM) tumor microenvironment (TME). Our findings establish, for the first time, that the oncolytic potential of ZIKV extends beyond direct cytolysis, encompassing a potent paracrine effect mediated by type III IFN that disrupts tumor-supporting vascular and immune networks both *in vitro* and *in vivo*.

The heterogeneity of GBM response to ZIKV categorized here into responsive and non-responsive phenotypes parallels clinical observations of viral therapy resistance. As previously demonstrated, responsive cells exhibit a viability reduction exceeding 50% (*34*). However, our data reveal a crucial insight: even in non-responsive cell lines, such as GBM02, which exhibit high viral titers without significant cell death, the establishment of viral replication serves as a signaling hub. This demonstrates that even if the tumor mass is not highly sensitive to ZIKV-induced apoptosis (*17*), infected cells can still release viral particles and signaling molecules that modulate the microenvironment and impact tumor progression.

A hallmark of GBM malignancy is its aberrant angiogenesis. We observed that factors secreted by ZIKV-infected GBM cells significantly impair endothelial migration, proliferation, and barrier integrity. While previous models have shown ZIKV-induced vascular defects during development (*35*), our study demonstrates that this property can be repurposed to starve the TME. The capacity of the ZIKV E protein to modulate endothelial function and interfere with tight junctions (*36*) further supports the concept that viral components act as multi-target inhibitors of tumor vascular stability. Within the clinical context of GBM, this controlled vascular disruption could potentially enhance the intratumoral delivery of systemic therapies.

Simultaneously, we observed a dramatic shift in the microglial landscape. The transition of microglia from a pro-tumoral state toward an antitumoral phenotype marked by increased iNOS is a pivotal finding. Nitric oxide is a crucial immunoregulatory molecule whose increased concentration is associated with apoptosis induction in GBM models (*37, 38, 39*). Furthermore, the reduced expression of ARG1 and IL-4 is significant, as these markers are linked to high tumor proliferation and poor prognosis (*40, 41, 42*). The concomitant increase in IL-6, which can trigger apoptosis pathways in GBM (*43*), and IL-1β, which exhibits antitumoral properties (*44*), suggests that ZIKV-induced factors effectively reprogram the immune niche. However, this inflammatory response must be precisely controlled to avoid chronic inflammation that could favor tumor growth (*45*).

Our transcriptomic and functional analyses identify type III IFN as the central orchestrator of these effects. While most GBM research has focused on type I and II IFN (*31, 32, 33*), type III IFN signaling offers a distinct translational advantage due to its potentially more localized action and reduced systemic inflammatory profile, stemming from the restricted expression of its receptor complex. Binding of IFNL1 to the IFNLR1/IL10RB complex activates JAK1/TYK2 signaling, driving the expression of interferon-stimulated genes (ISGs) such as *OAS1*, *MX1*, and *IRF9* (*46*). The upregulation of *SOCS1* observed in our data likely serves as a fine-tuning mechanism to limit survival signaling and promote programmed cell death (*47*). These findings align with reports that robust interferon signaling contributes to TME remodeling and enhanced T cell activation (*19, 48, 49*).

Our clinical validation using the CGGA database establishes a vital translational link, revealing that IFNLR1 and IL10RB expression is not only recapitulated in human GBM but peaks in the most aggressive and recurrent grades. This analysis exposes a critical paradox: patients with the most aggressive disease characterized by high IL10RB expression and maximal receptor density in recurrent tumors currently lack effective treatment options.

The significant correlation between high IL10RB expression and shorter overall survival identifies the type III IFN receptor complex as a hallmark of highly proliferative and treatment-resistant phenotypes. These high-risk patients possess the necessary molecular machinery to respond to type III IFN yet remain in a state of immune evasion due to a lack of endogenous ligands. This underscores an urgent unmet clinical need and presents a precision medicine opportunity to transform a marker of dismal prognosis into a therapeutic vulnerability, potentially rescuing survival in both primary and treatment-refractory glioblastoma.

Finally, we demonstrated that exogenous administration of type III IFN phenocopies the effects of ZIKV-conditioned media, significantly reducing tumor volume *in vivo*. This transition from a viral-based to a cytokine-based intervention is a critical translational step it leverages the potent TME-remodeling capacity of ZIKV while bypassing safety concerns associated with live neurotropic viral replication in patients. This aligns with emerging strategies where cytokines like IL-12 enhance CAR-T cell efficacy and remodel the TME (*50*). Similar to adenoviral vectors producing CXCL11 (*51*), our data reinforce that type III IFN expression is a potent therapeutic mechanism. Future studies should evaluate whether combining ZIKV-based oncolytic strategies with immune checkpoint blockade enhances antitumor efficacy by harnessing the microglial reprogramming described here. Moreover, defining the viral determinants, including nonstructural proteins, that drive type III interferon induction may facilitate the development of virus-derived therapeutic platforms while mitigating the neurotoxicity risks associated with live virus administration. Ultimately, these findings identify the type III interferon axis as a potent therapeutic vulnerability in glioblastoma, offering a novel strategy to overcome the immunosuppressive landscape of aggressive brain tumors.

## MATERIALS AND METHODS

### Study Design

The primary objective of this study was to characterize the oncolytic and immunomodulatory effects of Zika virus (ZIKV) on the glioblastoma (GBM) tumor microenvironment. We employed an integrative experimental approach combining in vitro assays with established cell lines (U-87 MG, TG1, GBM02, T98G) and an intracranial syngeneic mouse model using GL261 cells implanted in 12-week-old male Swiss mice. For in vivo experiments, mice were randomly assigned to treatment groups (MOCK vs. ZIKV) one-week post-inoculation, Intratumoral injections of 10^3^ PFU of ZIKV were performed using stereotaxic coordinates. In vitro mechanistic studies utilized conditioned media (CM) to isolate paracrine effects on endothelial cells and microglia, with all experiments performed in at least three independent biological replicates (n=3). To unravel the molecular mechanism, an unbiased RNA-Seq transcriptomic analysis was conducted, followed by functional validation of the identified type III interferon axis. Investigators were not blinded during data collection, but quantitative histological analyses (vessel density and microglial morphology) were performed using standardized automated software parameters to minimize bias.

### Ethics and Animal procedures

This study was approved by the Ethics Committee of the Center for Health Sciences at the Federal University of Rio de Janeiro (UFRJ) protocol N. A07/22-140-19. Swiss mice were obtained from and group-housed at five animals per cage at the Animal Facility of the Biomedical Sciences Institute at UFRJ. A total of 6 x 10^5^ cells of the murine glioma cell line GL261 was implanted into 12 week-old male mice. Mice anesthetized by intraperitoneal injection of Ketamine (100 mg/Kg) and Xylazine (25 mg/Kg) were placed in a stereotactic apparatus, and a brain midline incision was made on the scalp. The skull was punctured at stereotaxic coordinates: 0.5 mm posterior to the bregma and 2 mm lateral from the midline and 3,5 mm depth. Cells were delivered in 3µL with a 29.5 Hamilton syringe. One week after tumor implantation the animals were infected with 10^3^ PFU of ZIKV intratumorally. Matrigel (BD Biosciences, catalog number 354230) was mixed with conditioned media under CM MOCK and CM ZIKV conditions at a ratio of 24 volumes of Matrigel to 1 volume of conditioned media. Subsequently, 450 µL of this mixture was injected subcutaneously into the flanks of 8-weeks-old adult male mice. The plugs were retrieved one-week post-implantation, fixed in 4% paraformaldehyde, and embedded in paraffin. Serial sections of 5 µm thickness were obtained from each plug and stained with hematoxylin-eosin or CD31 to identify endothelial cells.

### BrdU

Cells were plate in 96-well plates and treated for 48 hr with GBM CM. Their proliferation capacities were determined by quantification of the BrdU incorporation into the DNA of replicating cells using the Cell Proliferation following the manufacturer’s instructions (Roche). After treatments, cells were incubated with BrdU labeling solution (0.1μL/mL) for 30 min or 1 h, respectively, at 37°C in a humidified atmosphere (5% CO2). Cells were then incubated with FixDenat solution and anti-BrdU POD (anti-BrdU-FLUOS) according to the manufacturer’s instructions. Colorimetric analyses were performed using a multi-label plate reader (Bio-Rad) and absorbances were determined at 450 nm.

### Cell culture

The U-87 MG, T98G, and GL261 cell lines were obtained from the Cell Bank of Rio de Janeiro (BCRJ). The GBM02 and TG1 cell lines, both patient-derived, were established and characterized following protocols (23,24) respectively. HBMEC was kindly provided by Professor Catarina de Moura Elias de Freitas. Microglial cultures were obtained from a pool of 10 cerebral cortex of newborn Swiss mice as previously described (*25*). Briefly, newborn mice (1-2 days after birth) were quickly decapitated, and the cerebral cortex dissected for use in primary cultures. Cells were plated on precoated poly-L-lysine plates, after 12 days in culture, floating microglial cells were isolated and plated for each experiment. All cells were cultured in Dulbecco’s modified Eagle’s medium (DMEM/F12) supplemented with 3.5 mg/ml glucose, 0.1 mg/ml penicillin, 0.14 mg/ml streptomycin and 10% fetal bovine serum (FBS). The cultured cells were maintained at 37°C in an atmosphere of 95% air and 5% CO2.

### Cell viability assay

Cell viability was assessed by Cell titer blue assay (Promega) according to manufacturer’s instructions. GBM cells were plated into 96-well plate for 24h after that cells were infected with the following MOIs 1, 3, 5 and 10 of ZIKV. Cell viability was measured after 3-, 5- and 7-days post infection (dpi). Cell titer blue was added in each well and incubated for 2 h at 37°C. After that, fluorescence was measured in a plate reader (Emax Plus, Molecular Devices) at 560nm excitation and 590nm emission.

### Conditioned medium preparation

1.8 x 10^5^ U-87 MG cells were plated per well in a 6-well plate. Cells were either MOCK or ZIKV-infected with an MOI 3 after 24h in culture. After 1h of adsorption, the inoculum was removed, and 2 mL of fresh medium supplemented with 10% FBS was added. The conditioned medium (CM) was collected 48 h after ZIKV infection and was filtered with an Amicon centrifugal filter (Milipore) with a 100 kDa filter to remove viral particles. The CM from the MOCK and ZIKV conditions underwent the same type of processing.

### ELISA assay for type III IFN

Type III IFN concentration in conditioned medium was assessed using IFNL1 Elisa kit (Invitrogen) according to manufacturer’s instructions.

### Gene expression analysis

The Ensemble of Gene Set Enrichment Analysis (EGSEA) package was used to identify classes of genes that could be underrepresented among the large set of differentially expressed genes and have some association with the oncolytic potential of ZIKV (26). The raw transcriptome data is uploaded to the Transcriptome Shotgun Assembly (TSA) repository, NCBI.

### Immunostaining

Cells were cultured on coverslips in a 24-well plate. After 48h cells were fixed with 4% paraformaldehyde for 20 min, permeabilized with 0.1% Triton X-100 in PBS, and blocked with 5% bovine serum albumin (BSA) for 1h. Cells were incubated overnight at 4°C with the following primary antibodies diluted in the blocking buffer: rabbit anti-GFAP 1:400 (Dako), mouse anti-4G2 1:1, rabbit anti-cleaved caspase 3 1:400 (Cell Signaling), rabbit anti-ZO1 1:400 (Invitrogen), rabbit anti-iNOS 1:400 (Millipore), isolectin-B4 1:400 (Invitrogen), rabbit anti-Iba1 1:10,000 (WAKO). Thereafter, the cells were washed with PBS and incubated with secondary antibody conjugated with Alexa Fluor 546, Alexa Fluor 488 or Alexa Fluor 647 (1:400) for 2h at room temperature. Cells were then washed with PBS, stained with DAPI, and mounted in Fluoromount-G®. After immunostaining, images were acquired at 63x with a laser scanning confocal SP5 microscope (Leica,). Data processing and analysis were conducted using ImageJ (National Institutes of Health).

### In Silico Analysis of Human Glioma Datasets

To assess the clinical relevance of the type III IFN receptor subunits (IFNLR1 and IL10RB) in human patients, we conducted a retrospective transcriptomic and clinical analysis using the Chinese Glioma Genome Atlas (CGGA) database. Database and Cohort Selection Transcriptomic data (mRNA-seq) and corresponding clinical metadata were retrieved from the mRNAseq_693 and mRNAseq_325 datasets. We evaluated the mRNA expression levels of IFNLR1 and IL10RB across distinct histological subtypes and malignancy grades according to the World Health Organization (WHO) classification. Samples were categorized into Oligodendroglioma (WHO II), Oligoastrocytoma (WHO II), Astrocytoma (WHO II), Anaplastic Oligodendroglioma (WHO III), Anaplastic Oligoastrocytoma (WHO III), Anaplastic Astrocytoma (WHO III), and Glioblastoma (GBM, WHO IV). Statistical differences between groups were determined using one-way analysis of variance (ANOVA) followed by Tukey’s post-hoc test for multiple comparisons. The prognostic value of the receptor subunits was evaluated specifically within the Glioblastoma (WHO IV) cohort. Patients were stratified into "High Expression" and "Low Expression" groups based on the median expression value (cutoff) of each gene. Overall survival (OS) was estimated using the Kaplan-Meier method, and the significance of survival differences between groups was assessed using the log-rank (Mantel-Cox) test. Hazard ratios (HR) and 95% confidence intervals (CI) were calculated to quantify the risk of mortality associated with receptor expression. Data visualization, including boxplots for expression distribution and Kaplan-Meier curves, was performed using R and GraphPad Prism.

### Microglial morphological analysis

The microglia image datasets from the in-vivo experiment were grouped and coded for unbiased analysis. Image processing was performed using Fiji (ImageJ). The entire microglia population identified from acquired images was analyzed to ensure reliability and statistical power. Morphological quantification of microglia cells included branching patterns and cell diameter measurements. Sholl Analysis was used to determine the Sum of Intersections (SI) (n = 371), while Feret’s diameter assessed cell size variations (n = 498). These methods helped differentiate between homeostatic and reactive states of microglia. The D’Agostino & Pearson Omnibus Normality Test assessed data normality, followed by the Mann-Whitney test for group comparisons. A p-value < 0.05 was considered statistically significant. Manual cell counting was performed while the data analysis in ImageJ. Student’s unpaired T-test was used to compare the average number of cells per image in both MOCK (n = 498) and ZIKV (n = 712) groups.

### Plaque assay

Viral titers were determined by performing a plaque assay on Vero Cells. Supernatant from infected (MOI 3) and mock cultures of GBM cell lines U-87 MG and GBM02 were serially diluted (10-fold dilution factor) and added to confluent Vero cell monolayers in 12-well plates. Cells were inoculated with the supernatant samples. After 1h, the inoculum was removed and semi solid medium (Alpha-MEM supplemented with 1% CMC and 1% FBS) was added. Cells were further incubated for 5 days and then fixed with 10% formaldehyde for 45 minutes at room temperature. Cells were stained with a solution of 1% crystal violet in 20% ethanol for 5 minutes and plaques were counted.

### qPCR

Microglia cells were maintained in culture until confluence state and treated with GBM conditioned media for 48 hours. After the treatment, total RNA was extracted using Trizol Extraction Reagent (Invitrogen) according to the manufacturer’s instructions. After this, cDNA was synthesized from total RNA using the High-Capacity cDNA Reverse Transcription Kit (Applied Biosystems).

Then, 500 ng of each cDNA sample was used for qPCR in a final volume of 25µL. For real-time fluorescence amplification and detection, SYBR Green (Applied Biosystems) was used as a probe with primers at a concentration of 0.05µM each and 40 cycles were performed, which the conditions of each cycle were: 15 seconds at 95℃ (denaturation) followed by 60 seconds at 60℃ (annealing and extension). The primer sequences used were ARG1 forward 5’-CTTGCGAGACGTAGACCCTG-3’ and reverse 5’-TCCATCACCTTGCCAATCCC-3’ ;β-actin: forward 5’-TGGATCGGTGGCTCCATCCTGG-3’and reverse 5’-GCAGCTCAGTAACAGTCCGCC-3’; IL-1β forward 5’-TACAAGGAGAACCAAGCAACGA-3’ and reverse 5’-TGCCGTCTTTCATTACACAGG-3’; IL-4 forward 5’-GAAGAACACCACAGAGAGTG-3’ and reverse 5’-AGCTCCATGAGAACACTAGA-3’; IL-6 forward 5’-GCCGAGTAGATCTCAAAGTG-3’ and reverse 5’-GCCGAGTAGATCTCAAAGTG-3’.

### Statical analysis

All values were expressed as mean ± SEM. Data normality was assessed using the Shapiro-Wilk test. For normally distributed data, comparisons between two groups were performed using Student’s t-test, and among multiple groups using one-way ANOVA followed by Dunnett’s post hoc test. For non-normally distributed data, appropriate non-parametric tests were applied. All statistical analyses were performed using GraphPad Prism 6 (GraphPad Software Inc). A p-value <0.05 was considered statistically significant.

### Transcriptome by RNA-seq

Total RNA isolation from GBM cells was done using TRIzol™ Reagent (Invitrogen™), followed by quantitation and quality analyses using Qubit RNA HS Assay Kit (Thermo Fisher Scientific, USA) and Agilent RNA 6000 Pico Kit (Agilent Technologies, USA). RNA-seq was performed using five biological replicates for each condition (MOCK and ZIKV) and the libraries were built using TruSeq Stranded Total RNA Library Prep Kits (Illumina, USA) according to manufacturer’s instructions. To evaluate fragment average and quantify libraries we used Agilent 2100 BioAnalyzer and High Sensitivity DNA Kit (Agilent Technologies, USA) and a qPCR-based KAPA library quantification kit (KAPA Biosystems, USA), finally we pooled libraries with 20 pM of final concentration to following sequencing steps. For paired-end sequencing we used the Illumina HiSeq 2500 Sequencing System (Illumina, USA) platform and HiSeq Paired-End Cluster Kits v4. The reads were aligned with STAR version 2.7.0f (Dobin et al., 2013) using the GENCODE project human genome version 30 (GRCh38.p12) as reference. The data were normalized using the TMM method (Trimmed Mean of M-values), using the Bioconductor edgeR package (Anders; Pyl; Huber, 2014). The selection of genes was made based on the corrected p values (FDR) ≤ 0.05 and fold change log (logFC) ≤ -1 or ≥ 1.

### Tube formation assay

Matrigel (BD Sciences) was added in 96-well plate and 30 minutes later the matrigel solidified and HBMEC cells were plated on top of the matrigel and were treated with GBM CM. Images were acquired at 2 and 8h after treatment in a phase-contrast Nikon Eclipse TE300 inverted microscope equipped with a digital camera (CoolSNAP-Procf color, ROPER SCIENTIFIC™ Photometrics).

### Viral detection by RT-qPCR

Viral RNA was extracted with the RNeasy Plus Mini Kit (QIAGEN, Venlo, The Netherlands), according to the manufacturer’s instructions. Viral load was determined by reverse transcription of RNA followed by quantitative PCR, performed with the GoTaq Probe 1-Step RT-qPCR System (Promega, Madison, WI, USA) on a 7500 Real-Time PCR System (Applied Biosystems, Foster City, CA, USA), using the primers and probe described by Lanciotti (27). ZIKV RNA copies were quantified by interpolation against a standard curve generated from eight 10-fold serial dilutions of a synthetic ZIKV RNA template corresponding to the targeted region.

### Viral infection

Cells were seeded in 6, 24 and 96-well plates for 24 h prior to infection. Cells were then infected with ZIKV-PE (GenBank: KX197192.1) at a multiplicity of infection (MOI) of 0.2, 1, 3 and 5 and incubated for 1 h at 37°C. After incubation, the inoculum was removed, and fresh medium was added, and cells were further incubated at 37°C with 5% CO_2_.

### Wound healing assay

Cell migration analyses through scratch assay was assessed through wound healing assay. A total of 2x10^5^ HBMEC cells were seeded on 24-well plates. After reaching 90% confluence, cells were treated with 10µM of cytosine arabinoside (Sigma Aldrich) overnight to prevent endothelial proliferation. After the treatment with GBM CM, a scratch was gently made with a 10µL pipette tip. Cell images were made at 0 and 24 hr and the cells that migrated to the risk area at 24 hr were quantified. Cells were observed in a phase-contrast Nikon Eclipse TE300 inverted microscope equipped with a digital camera (CoolSNAP-Procf color, ROPER SCIENTIFIC™ Photometrics).

## Acknowledgments

We thank all members of the Laboratory of Neuroplasticity, the Laboratory of Glial Cell Biology, and the Laboratory of Molecular Virology at the Federal University of Rio de Janeiro (UFRJ) for their invaluable discussions and insights throughout the development of this work. We are grateful to the Multiuser Confocal Microscopy Facility at the Institute of Biomedical Sciences (ICB/UFRJ) for technical support and infrastructure. We specifically thank Professor Catarina de Moura Elias de Freitas and Dr. Barbara Gomes da Rosa for kindly providing the HBMEC endothelial cell line. Additionally, we thank Fabio Jorge Moreira da Silva for essential assistance with animal care and maintenance at the animal facility.

## Funding

This work was supported by the Coordination for the Improvement of Higher Education Personnel (CAPES), the Carlos Chagas Filho Foundation for Research Support of Rio de Janeiro State (FAPERJ), and the National Council for Scientific and Technological Development (CNPq).

FAPERJ E-26/010.002165/2019

FAPERJ E-26/200.619/2022

FAPERJ E-26/201.365/2022

## Competing interests

The authors declare that they have no competing interests.

## Data and materials availability

The RNA-sequencing data generated in this study have been deposited in the NCBI BioProject database under accession number **PRJNA1412624**. All other data supporting the findings of this study are available within the article and its supplementary materials.

## Notes

https://www.ncbi.nlm.nih.gov/bioproject/?term=PRJNA1412624

### Competing Interest Statement

The authors have declared no competing interest.

