## Supplemental Material for "Zika virus oncolytic signaling reverts immunosuppressive and angiogenic glioblastoma microenvironment *in vitro* and *in vivo*"

**List of Supplementary Materials**

**Fig S1 to S3**

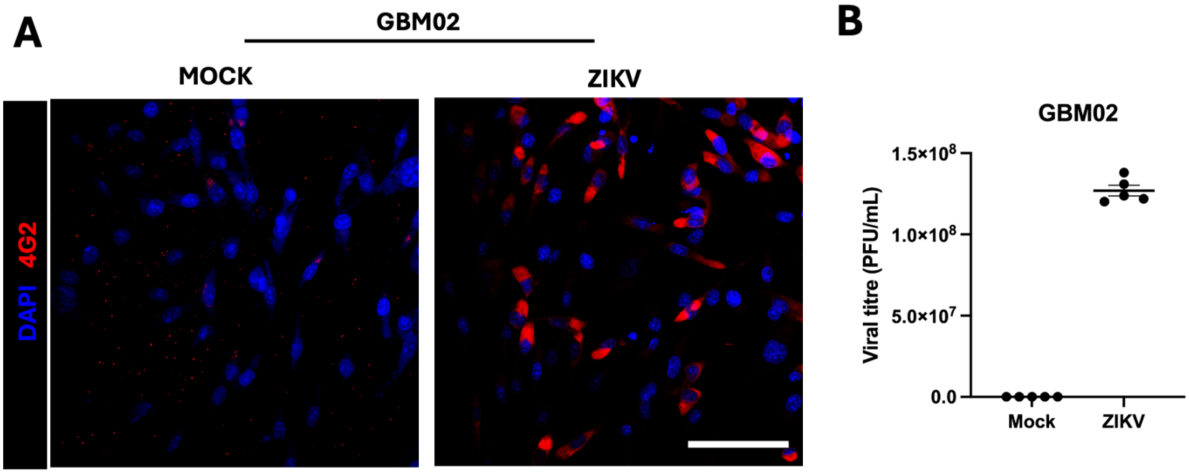


**Supplementary Figure 1. Viral production and infectivity in non-responsive GBM02 cells. (A)** Representative immunofluorescence images of GBM02 cells at 48 hours post-infection (MOI=3). Cells were stained for the viral envelope protein 4G2 (red) and nuclei (DAPI blue) demonstrating that approximately 90% of GBM02 cells are positive for the 4G2 marker, consistent with the high infection rates (96.7%) observed in the responsive U-87 MG phenotype. Data are presented as mean ± SEM of five independent experiments. **(B)** Viral titration of supernatants from GBM02 cells infected with ZIKV (MOI=3) 3dpi.

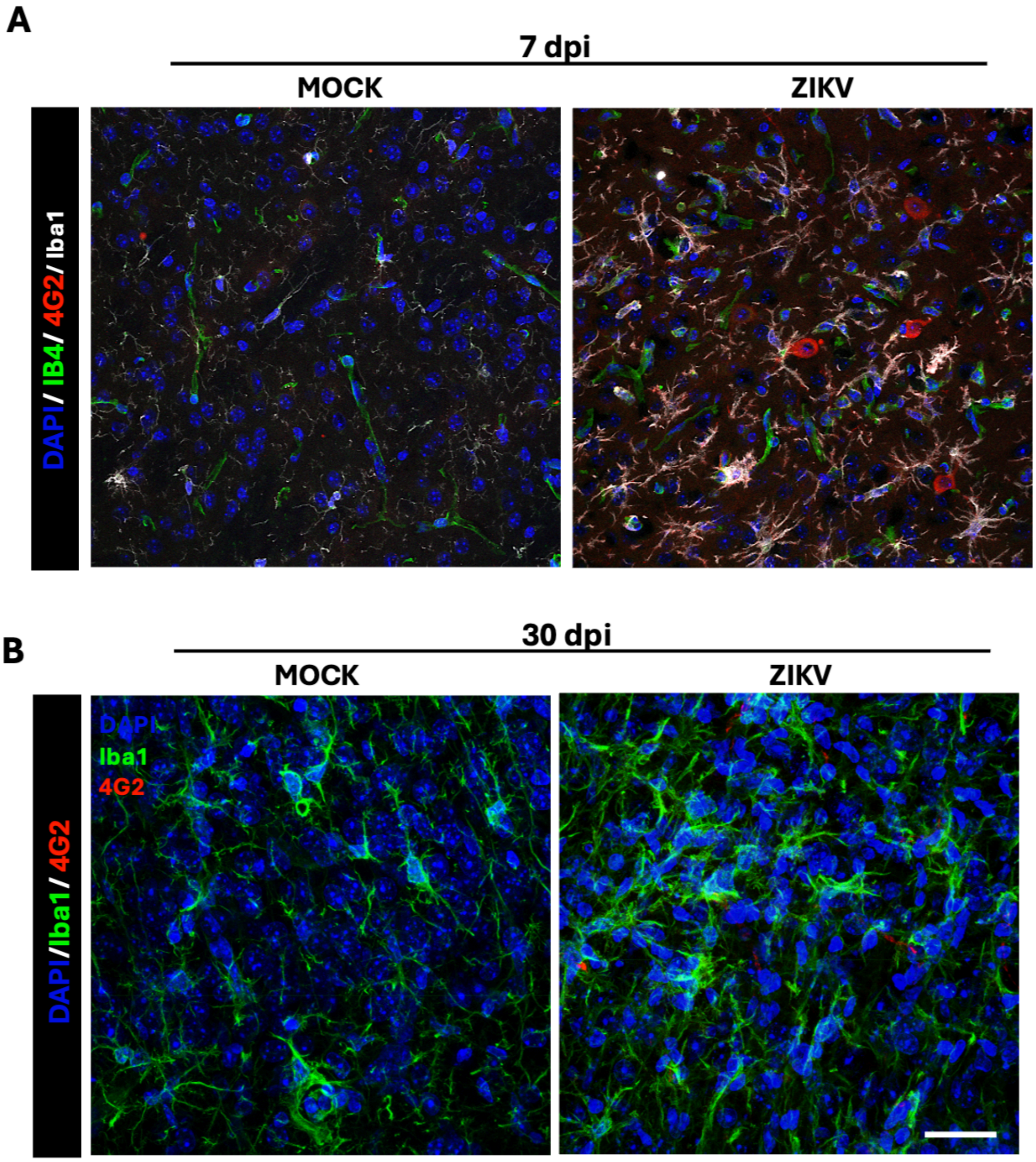


**Supplementary Figure 2: ZIKV persistence in GBM tissue *in vivo*.** Representative confocal laser scanning microscopy images of the tumor core from GBM-bearing mice. **(A)** Analysis at 7 days post-infection (dpi) and **(B)** 30 dpi, demonstrating the long-term persistence of the virus within the tumor mass. Sections are stained for nuclei (**DAPI**; blue), tumor-associated vasculature (**Isolectin B4/IB4**; green), microglial cells (**Iba1**; white), and the ZIKV envelope protein (**4G2**; red). Images were acquired at 40x magnification in the tumor core. Scale bars = 50 µm.

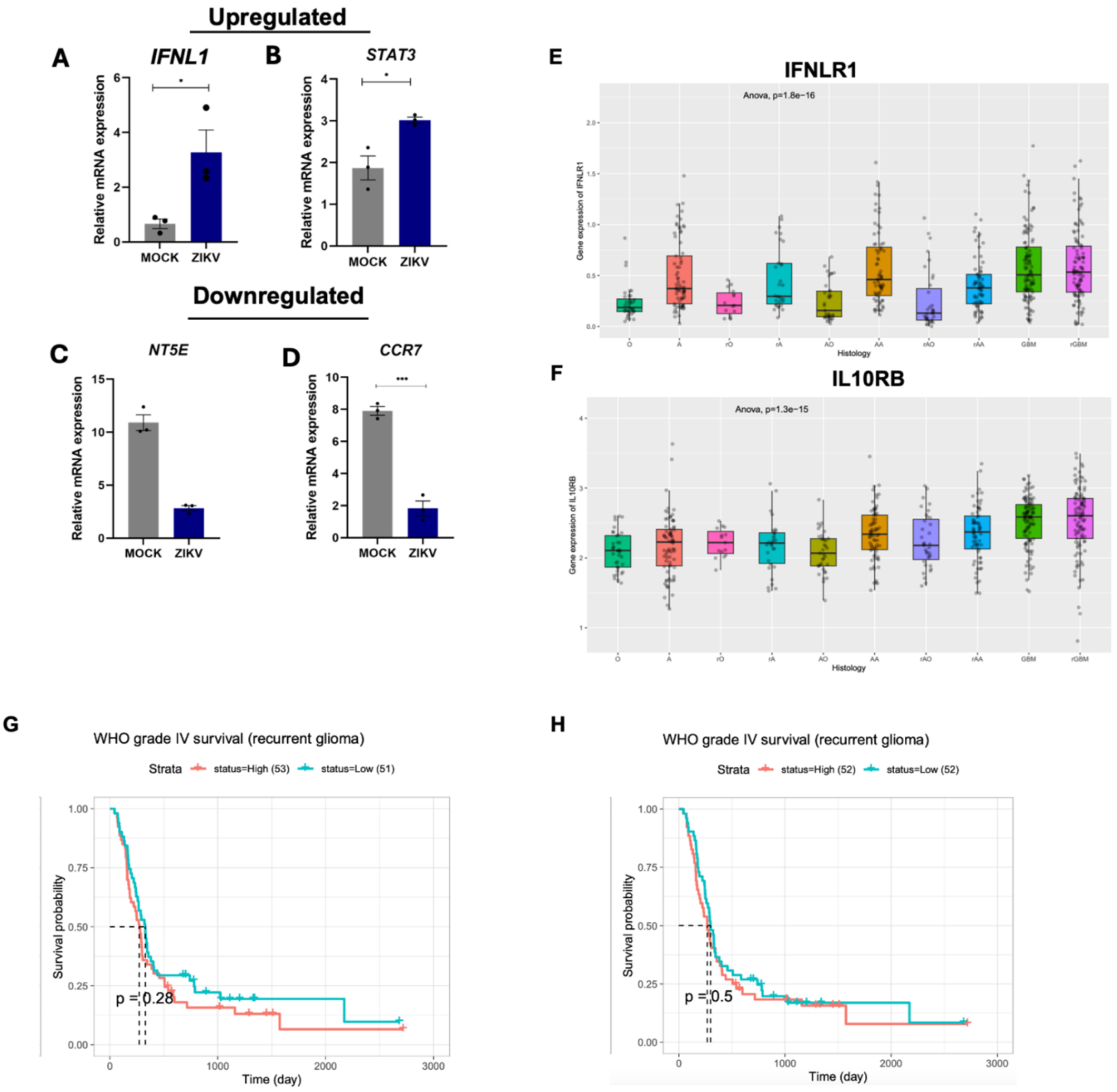


**Supplementary Figure S3. Validation of the ZIKV-induced Type III IFN signature and its clinical expression profile in glioma. (A–D)** Validation of transcripts in ZIKV-infected GBM cells. mRNA levels of *IFNL1*, *STAT3*, *NT5E* and *CCR7* detected in infected U-87 MG cells MOI=3 at 3 dpi. **(E–F)** Pan-glioma expression analysis of receptor subunits using the CGGA database**.** Boxplots demonstrate that *IFNLR1* and *IL10RB* mRNA levels are significantly elevated across various glioma subtypes, reaching peak expression in GBM samples. **(G–H)** Kaplan-Meier survival curves of patients with recurrent gliomas. Data in A–D are presented as mean ± SEM of three independent experiments (*$p < 0.05$, ***$p < 0.001$).
